# Drug repurposing for Leishmaniasis with Hyperbolic Graph Neural Networks

**DOI:** 10.1101/2023.02.11.528117

**Authors:** Yenson Lau, Jahir M. Gutierrez, Maksims Volkovs, Saba Zuberi

**Affiliations:** Layer 6 AI, Toronto, Ontario, Canada

## Abstract

Leishmanisis, a neglected tropical disease caused by protozoan parasites of the genus *Leishmania*, affects millions of individuals living in poverty across the world and is second to malaria in parasitic causes of death. Although current drugs for treating leishmaniasis exist, these are either highly toxic, ineffective, or expensive. For this reason, there is an urgent need to identify affordable, safer, and more effective treatments. Drug repurposing is a promising method for identifying existing molecules with the potential to treat leishmaniasis. Here, we present a deep learning model for drug repurposing based on hyperbolic graph neural networks. We leverage experimentally validated protein-drug interactions and molecular descriptors across three different parasites to train and validate our model. The final network model shows significant gains over the best baseline model, with an 11.6% increase in precision of the top scoring 0.5% protein-drug pairs. Finally, our model identified two experimental drugs that could target three *L. major* proteins involved in drug resistance and cell cycle regulation, which play an essential role in ensuring the parasite’s survival inside the host.

**Author summary:** Leishmanisis is a deadly and neglected tropical disease. Current treatments are ineffective, highly toxic, and unaffordable for the majority of affected individuals who live in extreme poverty. To accelerate the discovery and development of novel treatments against this disease, we develop a model that proposes existing drugs with the potential to treat leishmaniasis. Our methodology is based on hyperbolic graph neural networks, a class of deep learning models that can incorporate protein-drug interaction information from similar but well-studied parasites, in addition to utilizing chemical and molecular features. Empirically, this additional information leads to improved predictions of drug interactions with *L. major* proteins. Overall, the protein-drug pairs predicted by our model suggests two existing drugs that could target essential pathways for growth and survival of *Leishmania* parasites, which could be exploited either as a booster to increase the therapeutic effect of an existing anti-leishmaniasis drug, or as a novel chemotherapeutic treatment against the disease.

## Introduction

About 350 million people live in remote and poor areas with continuous exposure to vector sand flies (Phlebotomine) carrying one of many parasitic species of the genus *Leishmania*, which are responsible for causing the neglected tropical disease known as leishmaniasis. Depending on the type of leishmaniasis, the symptoms range from mucocutaneous ulcers to fatal visceral complications. Infected individuals may carry Leishmania parasites for long periods without any symptoms, but the disease eventually manifests clinically in the form of immunosuppression, fever, and anemia.

Today, many antiparasitic drugs and candidate vaccines exist as potential treatments against leishmaniasis [1]. Example drugs include the anticancer agent miltefosine as well as pentavalent antimony and pentamidine [2]. Unfortunately, these drugs are not fully effective as some Leishmania strains have developed resistance against them, and many of these drugs can cause serious adverse effects such as renal dysfunction, hepatitis, and cardiac arrhythmia. In addition, many of these treatments are unavailable in the countries where they are needed, and most are unaffordable for the majority of affected individuals.

To aid in the discovery and development of novel leishmaniasis treatments, many research groups have applied drug repurposing as a strategy to identify new uses for existing drugs. Drug repurposing leverages data from high-throughput screenings and clinical studies to train statistical and computational chemistry models that can recommend novel applications of existing drug molecules [3]. In fact, approximately 30% of all new drug approvals by the US Food and Drug Administration (FDA) in the last ten years originated from drug repurposing studies and more than 60% of current drugs for leishmaniasis (including the aforementioned drugs miltefosine and pentamidine) were repurposed from other indications [4]. Thus, because of the limited research and development resources allocated to neglected diseases like leishmaniasis, drug repurposing is an attractive and cost-effective method for developing new treatments, which can reduce the development process by 5 to 7 years [5–7].

Recent advances in high-throughput biology, pharmaceutical chemistry, and computer science have facilitated the creation of vast protein-drug interaction databases. These databases catalog millions of protein-drug pairs across multiple organisms and conditions. In this context, graph theory is an essential tool to analyze the complex networks that emerge from piecing together many protein-drug interactions [8, 9]. Furthermore, novel machine learning frameworks have been developed to specifically handle network data and make relevant predictions at the node or edge level [10, 11]. Given the rich biochemical data available for protein and drug molecules, machine learning algorithms such as graph neural networks [12, 13] (GNNs) benefit from combining molecular features with network interaction information to better learn node representations or embeddings [14–17]. Although powerful, many of these recent machine learning models were developed using data from non-parasitic human diseases and incorporate information from a single organism (e.g. mouse or human) [18–20]. This limits their application in neglected tropical diseases where the target proteins to be drugged are not of human origin and instead belong to the target parasite’s cells. Furthermore, modern GNN architectures that exploit richer node representations with the use of hyperbolic geometry [21, 22] have been shown to outperform state-of-the-art GNNs across a variety of tasks [23], but have not yet been applied in drug repurposing.

Here, we propose a new approach, Hyperbolic Graph Neural Networks for Drug Repurposing (HGNN-DR). This is, to the best of our knowledge, the first drug repurposing model for leishmaniasis based on hyperbolic graph neural networks (HGNNs). Our model leverages experimentally validated protein-drug interactions across *Leishmania major* and two other eukaryotic parasites: *Plasmodium falciparum*, and *Trypanosoma brucei* and provides an easily extendable framework to include additional organisms. For each node in our network (drug or protein), we compute molecular attributes which HGNN-DR uses as input features and combines with network information for the task of predicting new protein-drug interactions (i.e. binding).

Our main contributions can be summarized as follows. First, we integrate protein-drug interaction data from *multiple* parasites in a novel way and leverage these experimentally validated interactions to better recommend drugs for targeting *L. major* proteins. This not only allows for building a larger training dataset but it also allows the model to utilize information about candidate drugs contained in the interaction networks of other parasites to learn better representations of these drugs. We demonstrate this by comparing our final model against neural networks trained on single-organism datasets. Second, we show that machine learning models that embed protein-drug interaction networks using hyperbolic geometry are advantageous for drug repurposing, by comparing our HGNN-DR model against standard GNN architectures in Euclidean space. Furthermore we demonstrate the performance gain of our approach over machine learning models that rely only on protein and drug features. Finally, we use our model to predict protein-drug interactions that could be exploited either as a booster to increase the therapeutic effect of an existing anti-leishmaniasis drug, or as a novel chemotherapeutic treatment against the disease.

## Methods

### Overview

Our drug-recommendation pipeline consists of three major components (Figure 1A-C). First, we construct a network consisting of known protein-drug interactions for *L major* as well as two supporting organisms, *P. falciparum* and *T. brucei*, from the STITCH database [24]. Next, we extract features for each drug using the Mordred and RDKit chemoinformatics toolkits [25, 26], and extract UniRep features for each protein sequence [27]. Finally, we train our HGNN-DR model on the known interaction network, combined with the corresponding protein and drug features, to predict the likelihood of *unknown* protein-drug interactions for drugs never tested experimentally against *L. major*. We compare the hyperbolic graph neural network against a standard Euclidean GNN approach, as well as feature-based models that predict the likelihood of a protein-drug interaction based only on the corresponding features.

**Fig 1.**
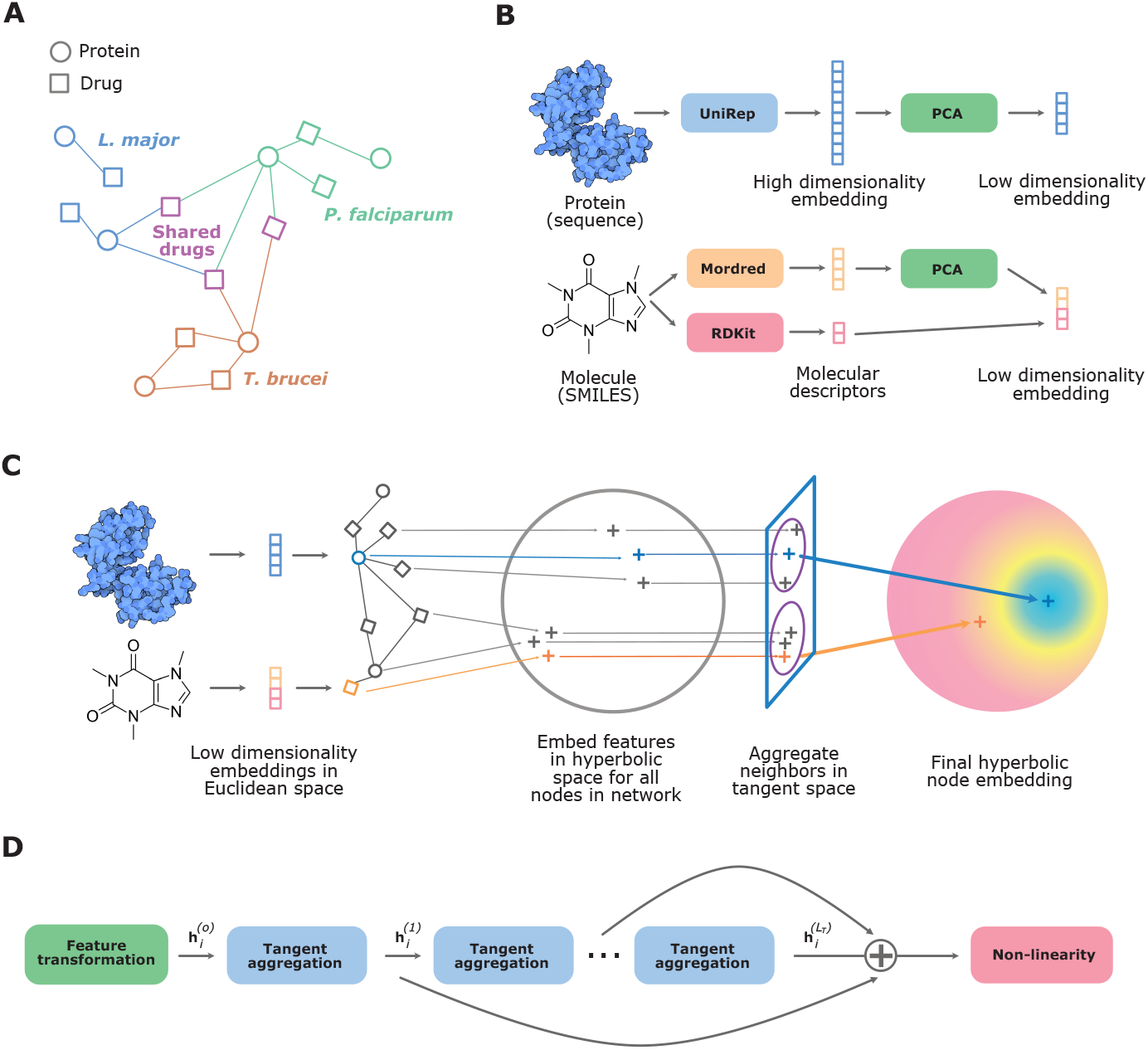
Hyperbolic embedding construction. (A) Simplified representation of the multi-organism protein-drug interaction network. (B) High-dimensional embeddings in Euclidean space are computed for proteins using UniRep and for drug molecules using a combination of Mordred and RDKit. The dimensionality of these embeddings is reduced via principal component analysis. (C) For all nodes in the protein-drug interaction network, the low-dimensionality Euclidean embedding of the node and the information of its location in the network are combined to create an embedding in hyperbolic space. By aggregating the neighbor nodes in the tangent space, a final hyperbolic node embedding is computed. (D) Each layer of the HGNN-DR encoder consists of a feature transformation, multiple layers of tangent aggregation with a residual connection and a non-linearity.

### Extracting Protein-Drug Interactions

The protein-drug interaction network was built using data from the STITCH database [24]. We downloaded the full protein-chemical interaction networks of *Leishmania major* (taxid 5664), *Plasmodium falciparum* (taxid 5833), and *Trypanosoma brucei* (taxid 5691). We select the latter two parasites due to their taxonomic proximity to *L. major* and the similarity of their infection processes and responses to drug treatments, making these useful drug repurposing signals for *L. major*.

#### Filtering

To create a high quality protein-drug network, we filtered the original interaction data from STITCH as follows: (1) We only keep protein-chemical pairs with experimental evidence, and discard interactions with weaker levels of confidence derived from text mining or computational predictions. (2) We only keep interactions involving stereo-specific compounds. Compounds with “flat” structures (merged stereo-isomers) are discarded. (3) We identify drugs by matching STITCH chemicals to drug identifiers in either the KEGG [28] or DrugBank [29] databases; any unmatched chemicals are discarded. (4) Finally, we discard any protein-drug interactions flagged as “catalysis” to exclude promiscuous enzymes involved in metabolism of xenobiotics. After filtering, the remaining interactions form a bipartite network, with nodes representing proteins and drugs and with edges representing the interactions between them.

#### Known Anti-Leishmaniasis drugs

To further evaluate our models, we included two known anti-leishmaniasis drugs, miltefosine and pentamidine [2], to ensure they are present in the test dataset. Interestingly, the interactions of miltefosine with *L. major* proteins are annotated in STITCH at a level below experimental evidence, thus miltefosine is discarded from our protein-drug interaction network during the data filtering steps described above. Nevertheless, we included these miltefosine-protein pairs in our test set given the actual therapeutic usage of this drug in treating leishmaniasis.We therefore included non-experimental interactions for miltefosine and filter them using steps (2)-(4) described above, before adding them to the test set, the construction of which is described in detail in Data Splitting.

### Extracting Protein and Drug Features

We used two chemoinformatics libraries, RDKit [26] and Mordred [25], to extract 199 and 1,375 numerical descriptors for drugs, respectively. We used UniRep [27] to extract features in the form of 1,900-dimensional vectors for proteins. After filtering and dimensionality reduction, features for drugs and proteins are represented by 240 and 100 dimensional vectors, respectively. For further details, see Protein and Drug Features in the Appendix.

### Data Splitting

We split the protein-drug interactions of all three parasites, into the following sets: a training set for modeling, a validation set used for model selection, and a test set to report final performance. Our goal is to use the model to infer new interactions between the *L. major* proteins and drugs that are present in the *P. falciparum* and *T. brucei* networks, but have no known interactions with *L. major*. We refer to this set of unknown interactions as the inference set. A schematic of this data split is shown in Figure 2.

**Fig 2.**
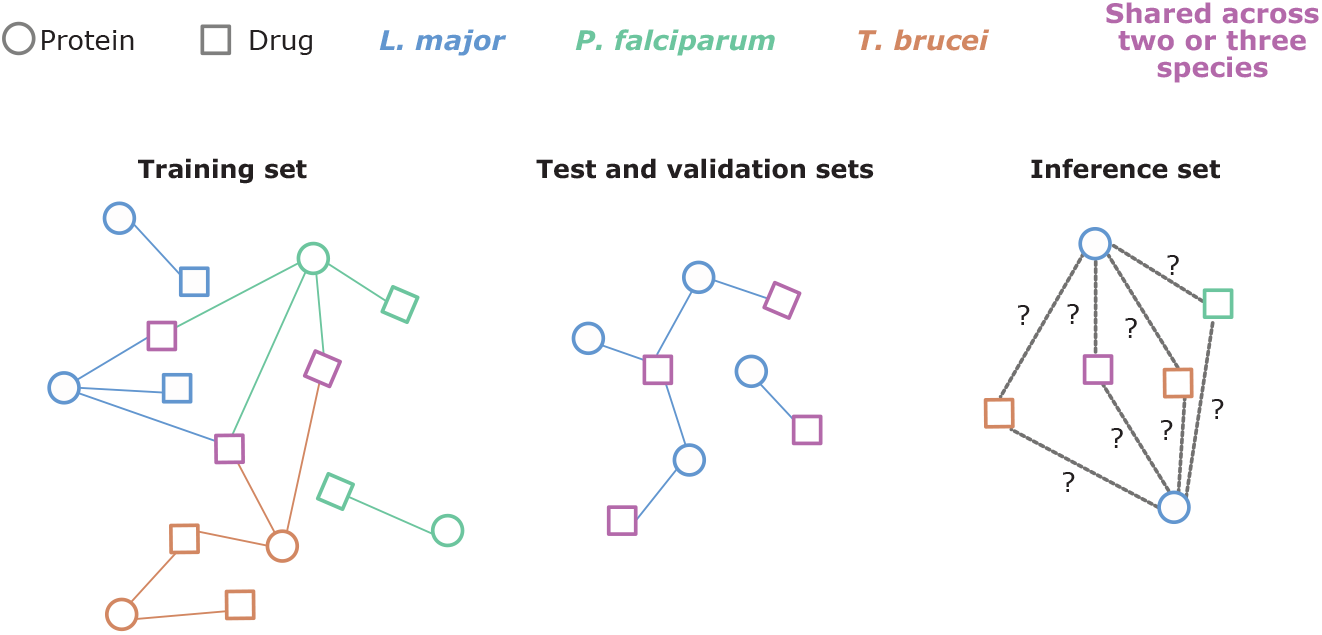
Data splitting schematic. Three datasets are used for training and validation. The training set contains protein-drug interactions from all three parasite species. The test and validation sets only include protein-drug pairs with a *L. major* protein (blue circles) and drugs found in *L. major* and at least one other species (pink squares). Finally, the inference set includes unknown interactions between *L. major* proteins and drugs found in either the *Plasmodium falciparum* or *Trypanosoma brucei* networks (green and orange squares, respectively).

To define these sets, we need to take into account the non-i.i.d. nature of our data; we expect the presence of interactions to depend on the specific protein or drug involved. Let *P***_L_** and *D***_L_** be the proteins and drugs in the *L. major* network, and *P***_PT_** and *D***_PT_** be the proteins and drugs from the union of *P. falciparum* and *T. brucei* networks. The *inference set* which, as described above, is used to provide drug recommendations for *L. major*, consists of unknown interactions from the set *P***_L_** × (*D***_PT_** *\ D***_L_**). Next, the drugs in *D***_L_** *∩ D***_PT_** (i.e. drugs which are present in the *L. major* network and at least one additional parasite network) are partitioned into a test and validation drug sets *D*_test_ and *D*_val_ The interactions in *P***_L_** × *D*_test_ and *P***_L_** × *D*_val_ are then used as the *test and validation sets* to evaluate the model’s ability to transfer information about drug interactions from other parasites to *L. major*. Finally, the remaining interactions are used for *training only*. A detailed figure of the data splitting is provided in Fig S1.

### Hyperbolic Graph Neural Network for Drug Repurposing

Here we present HGNN-DR, a hyperbolic graph neural network for drug repurposing, that predicts the likelihood of interaction between the drugs and proteins in our network. The model embeds features in to hyperbolic space and uses an encoder to learns *n*-dimensional protein and drug representations. The HGNN-DR encoder layer applies a feature transformation and multiple sub-layers of neighbourhood aggregation with residual connections (see Figure 1D). To predict the likelihood of interaction between a given drug and protein, we use a Fermi-Dirac decoder [21] based on hyperbolic distance, where the closer the protein and drug representations are in hyperbolic space, the more likely they are to interact.

A critical aspect of GNNs is neighbourhood aggregation, where node representations are learned by iteratively aggregating information from 1-hop neighbours in the network. In this way protein features are updated with information from neighbouring drugs and visa-versa. However, prior work has shown that GNNs are often unable to aggregate over multiple steps in the network without degrading performance [30]. Our approach extends prior HGNNs [21, 22] to address this issue. We decouple feature transformation and neighbourhood aggregation, allowing for multiple hops of neighbourhood aggregation for each layer of feature transformation, and further utilize residual connections [31–33] to deal with over smoothing. We show that HGNN-DR is better able to aggregate information over multiple hops in the network compared to the standard HGNN approach. Below we introduce some useful notation, and then describe the components of our HGNN model. We present additional background on hyperbolic space relevant to our model in the Appendix.

#### Notation

Given *s* proteins *P* = {*p*_1_, *…, p_s_*} and *t* drugs *D* = {*d*_1_, *…, d_t_*}, the known interactions between proteins and drugs are given by a sparse binary *s* × *t* matrix *R*, where *R_pd_* = 1 indicates an interaction. We denote the set of drugs a protein *p* interacts with as *𝒩_p_* = {*d* ∈ *D* | *R*_*pd*_ = 1}, which represents the neighbourhood of node *p* in the protein-drug graph, and similarly 𝒩*_d_* = *p P R_pd_* = 1 denotes the set of proteins that the drug *d* interacts with.

#### Embedding Features in Hyperbolic Space

The input features are extracted as described in Extracting Protein and Drug Features, yielding 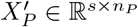 and 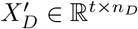*\* Euclidean inputs for proteins and drugs, respectively. These inputs are combined to yield a feature matrix 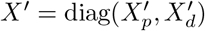.

For a point **x** ∈ *ℬ^d^* in hyperbolic space, its tangent space *𝒯***_x_***ℬ^d^* is a Euclidean space that approximates *ℬ^d^* at **x**. The exponential map exp**_x_** : *𝒯***_x_***ℬ^d^ →ℬ^d^* and logarithmic map log**_x_** : *ℬ^d^ → 𝒯***_x_** *ℬ^d^* allow points in the tangent space to be mapped to hyperbolic space and visa-versa and are defined in the Appendix. We assume the Euclidean features for each node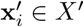 belong to the tangent space at the origin **o** of the Poincaré Ball. They are mapped to hyperbolic space via the exponential map, 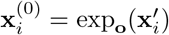

#### Encoder Layer

The HGNN-DR encoder layer includes feature transformation and *multiple* sub-layers of neighbourhood aggregation. An overview is shown in Figure 1D. Hyperbolic feature transformation involves multiplication by a weight matrix, *W* ∈ *ℝ*^n × *n*^, and the addition of a bias term, **b**′, defined on the tangent space *𝒯***_o_***ℬ*, following the approach in [34]:

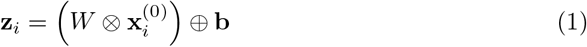

where **b** = exp**_o_**(**b**′), and the operators ⊗ and ⊕ are defined in the Appendix.

In our formulation, aggregation is carried out on the tangent space at **o** and so the log map, 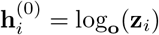 is used to project the representations **z***_i_* from *ℬ* to *𝒯***_o_***ℬ*, prior to aggregation. For node *i*, with neighbourhood *𝒯_i_*, the representation is updated as:

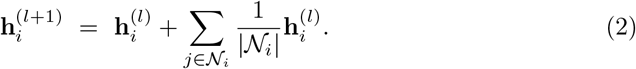

The residual connection then combines the representations from all intermediate layer 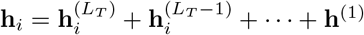, where *L_T_* is the number of layers of tangent space aggregation. Lastly we apply a non-linearity in Euclidean space, *σ*, before projecting back to hyperbolic space using an exponential map and obtaining our output node representation, **x***_i_*, as

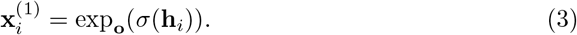

#### Decoder

To compute the likelihood of a link between a given protein and chemical node, we use the Fermi-Dirac decoder [21, 23], which utilizes the distance, *d_ℬ_*, between protein-chemical node representations computed in hyperbolic space (see Appendix)

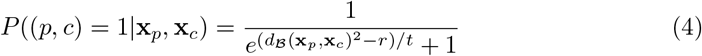

with hyper-parameters *r, t*.

### Model Evaluation

#### Baseline Models

We compare the performance of HGNN-DR to the following models:

- *Logistic regression*, one of the simplest learned classifiers.
- Two *gradient-boosted decision trees*, XGBoost [35] and LightGBM [36], which are amongst the strongest performing models for tabular data.
- A *multi-layer perceptron* (MLP), a neural network that performs several layers of feature transformations to the input before feeding the final output into a logistic regression classifier.
- *Euclidean graph neural network* (EGNN) [12], which performs layer-wise feature transformations and neighbourhood aggregation.

The first three baselines are feature-based models that do not explicitly incorporate interaction information. Instead, predictions of interactions between a drug-protein pair is based on their features only. While the EGNN includes network information through aggregating features over neighbouring nodes in the graph, similar to HGNN-DR, but without embedding node representations in to hyperbolic spaces.

#### Evaluation Metrics

Each model is expected to give high prediction scores for likely protein-drug interaction pairs. We measure model performance via several metrics calculated based on the model scores on the test set:

- Area under the receiver operator characteristic curve (AUCROC) and area under the precision-recall curve (AUCPR).
- Top-*q*% global, and top-*k* local precision and recall. Here, top-*q*% global metrics measure performance on the *q*-th percentile highest scoring interaction pairs overall. Top-*k* local metrics measure the performance on the *k* highest scoring interactions for each protein, averaged across all proteins.

AUCROC and AUCPR are common metrics used for summarizing overall classification performance. Global metrics, which look for top ranked interactions across all proteins and drugs, are useful for finding candidates for drug discovery. Finally, local metrics are useful for finding likely drug interactions given a particular protein for analysis.

## Results

### Protein-Drug Interaction Dataset

The size of the protein-drug interaction networks for *L. major, P. falciparum* and *T. brucei*, as well as the combined network, are summarized in Table 1. We see the interactions with *P. falciparum* and *T. brucei* contributes 57% of the edges in our combined network, and that there is a large overlap in the drugs present in these networks and the *L. major* network. This increases the training data relevant to *L. major* significantly.

**Table 1.** Protein-drug interactions from STITCH database, including links from known drugs.

| ORGANISM (TAX. ID) | # PROTEINS | # DRUGS | # EDGES |
| --- | --- | --- | --- |
| Leishmania major (5664) | 2,187 | 1,914 | 23,602 |
| Trypanosoma brucei (5691) | 1,947 | 1,817 | 20,224 |
| Plasmodium falciparum (5833) | 1,280 | 1,457 | 11,444 |
| Combined network | 5,414 | 2,065 | 55,270 |

As described in Data Splitting, we split our protein-drug interactions into training, validation, testing, and inference data. Since we wish our model to be able to transfer information about drug interactions from other parasites to *L. major*, each split is determined by the drugs involved in the protein-drug interaction such that drugs in the training split do not appear in validation, and so forth. While the training set includes edges involving proteins across all 3 parasites, the validation, testing, and inference edges involve only the 2,187 proteins belonging to *L. major*; all splitting here is done drug-wise. There are 151 drugs present in the *T. brucei* and *P, falciparum* networks that have no known edges in the *L. major* network, and these make up the inference set.

The drugs for the test set are selected such that approximately 10% of edges in the network are included in the test set. We perform 3-fold cross validation for hyper-parameter tuning and model selection.

### Model Performance

A summary of model test performance is shown in Figure 3. HGNN-DR significantly outperforms all other models in AUCPR, and consistently shows strong performance across the local and global precision and recall metrics. Although AUCROC is a common metric for evaluating classification tasks, it is not particularly useful here due to high label imbalance. AUCPR is more useful for model evaluation and comparison in such settings [37] and so we focus on this metric, along with the local and global precision and recall. Further performance metrics are reported in Table S1. Also see the Supplementary Material for Model Hyperparameters.

**Fig 3.**
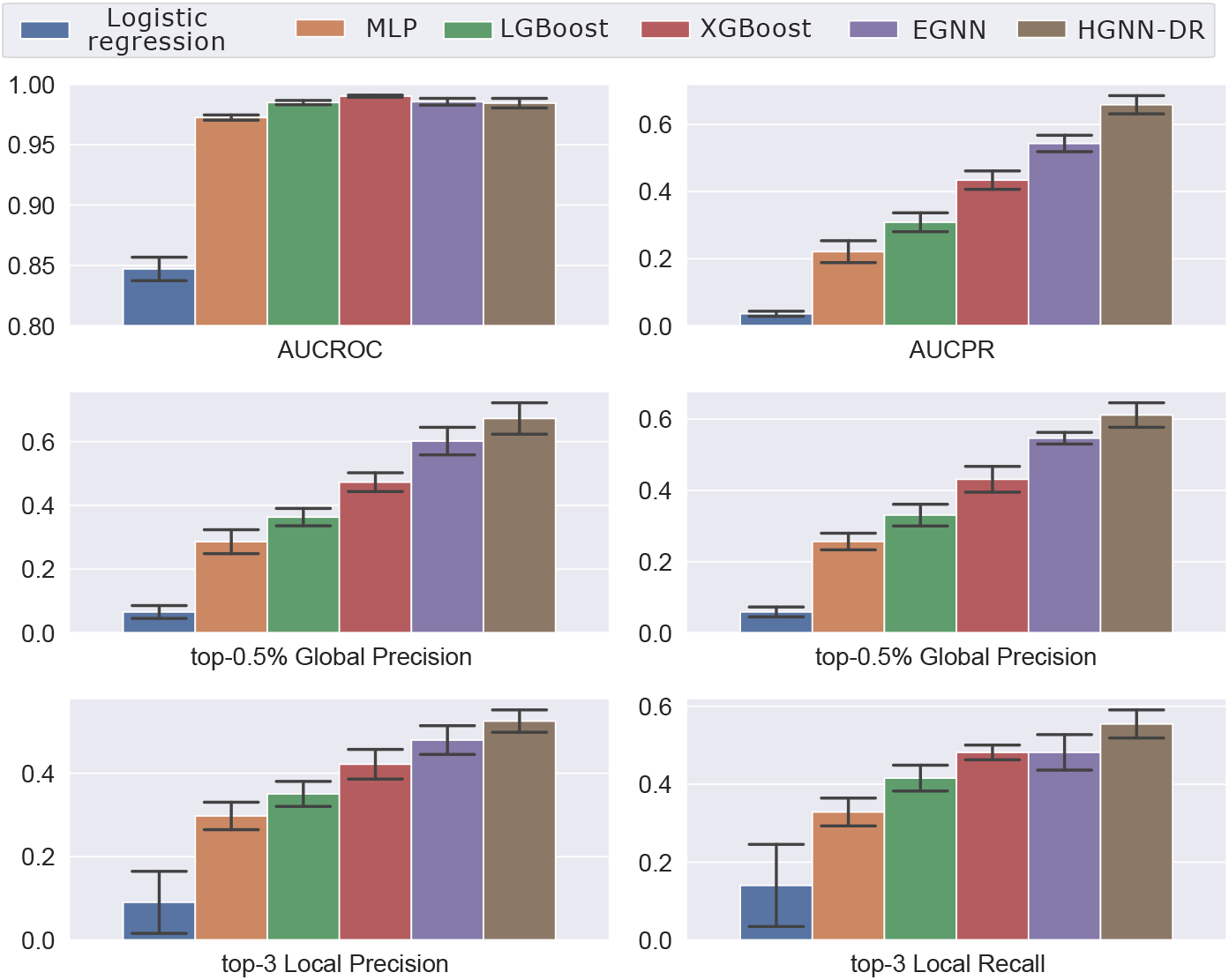
Comparison of model performance on the test set, of HGNN-DR against baseline models. Metrics shown are area under the receiver operating characteristic (AUCROC) and precision-recall (AUCPR) curves, top-0.5% global precision and recall, and top-3 local precision and recall. The bar height and errors represent the mean and standard deviation of performance evaluations over ten different random data splits.

We find that modeling protein-drug interactions directly using a graph neural network leads to significant performance improvements over feature based models. Our results show that the EGNN is able to outperform the feature-based models across local and global metrics. Modeling interactions in hyperbolic rather than Euclidean space with HGNN-DR gives further performance gains.

Amongst feature-based models, XGBoost provides the strongest performance, and both tree models perform much better than MLP or logistic regression. Although the MLP under-performs tree models, its model architecture is analogous to GNNs without modeling graph structure, making it a useful point of comparison. When compared to the MLP, we observe that HGNN-DR achieves large gains across all metrics.

Our HGNN-DR model modifies prior approaches, by decoupling the feature transformation and neighbourhood aggregation steps, as described in Hyperbolic Graph Neural Network for Drug Repurposing. In Table 2, we compare the performance of the standard HGNN [21] to our model with 2 layers of neighbourhood aggregation. While the standard HGNN performance drops at 2 layers, likely due to over-smoothing, our model is able to take advantage of longer range structure in the network. This demonstrates that the extra feature transformation and non-linearity in the standard HGNN is not only unnecessary but hurts the model performance.

**Table 2.** Ablation results comparing our HGNN model to prior approach from [21].

| Method | AUCPR | Top-0.5% Global Prec. | Top-0.5% Global Rec. | Top-3 Local Prec. | Top-3 Local Rec. |
| --- | --- | --- | --- | --- | --- |
| Standard HGCN 1 Layer | 0.570±0.027 | 0.587±0.043 | 0.534±0.030 | 0.486±0.032 | 0.512±0.049 |
| Standard HGCN 2 Layer | 0.477±0.042 | 0.545±0.053 | 0.495±0.034 | 0.453±0.034 | 0.464±0.050 |
| HGNN-DR | 0.659±0.029 | 0.672±0.052 | 0.612±0.036 | 0.525±0.028 | 0.555±0.038 |

#### Performance on Known Anti-Leishmaniasis Drugs

We further examine the ability of HGNN-DR to separate interactions from non-interactions on miltefosine and pentamidine, two known anti-leishmaniasis drugs. We compare the score distribution of known (or positive) interactions versus unknown (or negative) interactions to evaluate their separation, using the Kruskal-Wallis *H* and Mann-Whitney *U* tests.

For miltefosine, the two distributions yield test statistics of *H* = 13.70, with a × *p*-value of *p* = 2.15 × 10^−4^, as well as *U* = 10, 675; *p* = 2.15 × 10^−4^. The pentamidine × score distributions yields *H* = 5.46; *p* = 1.95 × 10^−2^ and *U* = 19, 433; *p* = 1.95 × 10^−2^. These results show that the HGNN-DR score distributions for positive and negative interactions are clearly distinct for known anti-leishmaniasis drugs, illustrating the discriminative power achieved by our model.

### Utility of Interactions from Additional Organisms

The addition of *P. falciparum* and *T. brucei* to the protein-drug interaction network is intended to provide two benefits: firstly, it provides previously unseen drug candidates for repurposing that may be likely to have interactions with proteins belonging the *L. major* network. Next, additional interaction information from these organisms may help to improve model performance for making predictions for such interactions.

Figure 4 summarizes the AUCPR achieved by our MLP, XGBoost, and HGNN models when the data included for training consists of a) *L. major* (tax. ID 5664) only; b) *L. major* and *P. falciparum* (tax. ID 5664, 5833); c) *L. major* and *T. brucei* (tax. ID 5664, 5691); and finally d) *L. major, T. brucei*, and *P. falciparum* (tax. ID 5664, 5691, 5833). We have found that the performance trends exhibited by the AUCPR metric are indicative of the trends exhibited by our other metrics.

**Fig 4.**
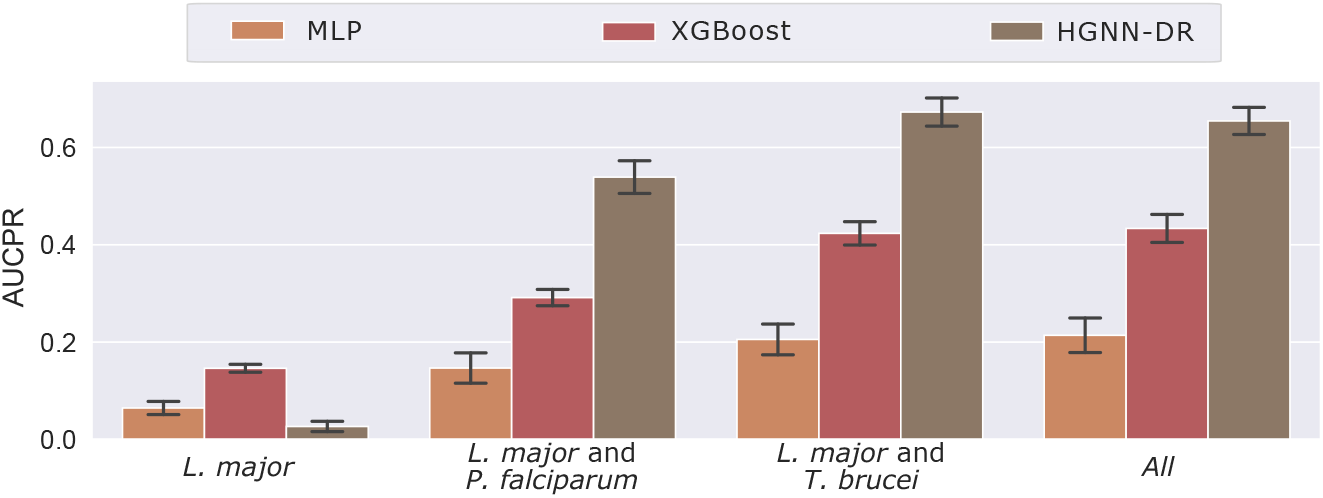
AUCPR gain by adding *P. falciparum* and *T. brucei*. Performance seen by MLP, XGBoost, and HGNN models with training data from *L. major* only, compared to adding *P. falciparum, T. brucei*, or both organisms to the training split. The bar height and errors bars represent the mean and standard deviation of performance evaluations over ten different random data splits / model fits, respectively.

It is clear that all models exhibit significant gains in performance when at least one additional organism is used to augment the *L. major* training data. In particular, our HGNN model, which relies heavily on modeling additional interaction information, underperforms feature-based models using *L. major* interactions alone. The addition of *T. brucei* provides a significantly larger performance gain compared to *P. falciparum* for all models. This is consistent with the fact that *T. brucei* is taxonomically closer to *L. major* than *P. falciparum*, and provides protein-drug interactions with closer overlap to *L. major*, both in terms of the proteins of the organisms and the overlap of drugs in their networks.

Furthermore adding *P. falciparum* after *T. brucei* does not provide a significant overall gain to performance on the test set. This is likely due to the lack of exclusive drugs added by *P. falciparum* to the test set. Although close to 70% of the drugs in the test set also exit in the *P. falciparum* network, only approximately 1% are exclusive to this organism. As a result, much of the interaction information for the drugs from *P. falciparum* would also have coverage from the *T. brucei* network, due to the heavy overlap of the drugs in the *L. major* and *T. brucei* networks.

Importantly, however, the *P. falciparum* network provides a large number of drugs to the inference set for drug candidate recommendation. Out of the 151 drugs in the inference split, 132 are exclusively provided by the *P. falciparum* network.

### Drug Recommendations

To emulate drug re-purposing recommendations with our final model, we ranked the protein-drug pairs in the inference set by their predicted score and looked at the top 50 scored pairs. We found that these pairs include 50 unique proteins from *L. major* interacting with only six distinct drugs. Most of these 50 proteins are annotated as transporters or putative protein kinases, and we identified two interesting protein-drug pairs.

First, the experimental drug Nu6027 (PubChem identifier 398148) was predicted to interact with two protein kinases involved in cell division (CRK1 and CRK3). Nu6027 is an aminopyrimidine that was found to inhibit cell growth in tumor cells by competitively inhibiting the human cyclin dependent kinases 1 and 2 (CDK1, CDK2) [38]. The 3D structure of Nu6027 bound to CDK2 is currently available on the Protein Data Bank with the identifier 1E1X.

Second, the experimental drug verlukast (PubChem identifier 6509849) was predicted to interact with a pentamidine-resistant protein coded by the gene PRP1. Verlukast is a potent antagonist of the cysteinyl leukotrine receptor 1 (CysLTR1) and it was once investigated as a candidate to treat chronic obstructive asthma. Interestingly, verlukast is in the top 5% most similar molecules to pentamidine in hyperbolic embedding space. This prediction suggests that verlukast may have a synergistic effect with pentamidine by blocking the anti-pentamidine resistance conferred by PRP1.

To better understand how our model predicted such protein-drug pairs, we looked for proteins in the training set with high similarity to PRP1, CRK1, or CRK3 and which were known to interact with either verlukast or Nu6027. First, we computed the hyperbolic distance between the embeddings of all proteins in the training set and the embeddings of these three proteins of interest. We found that several Trypanosoma and Plasmodium proteins were both close to these proteins of interest in embedding space and were known to interact with verlukast or Nu6027. In fact, after ranking the proteins in the training set by their proximity to PRP1, CRK1, or CRK3, we found that those proteins known to interact with verlukast or Nu6027 were among the top 0.5% closest proteins in embedding space (Figure 5), even when these proteins have a sequence similarity of less than 35%. All other *L. major* proteins predicted to interact with verlukast or Nu6027 were members of either the ATP-binding or ABC transporter protein subfamilies, which is expected given the versatility of membrane transporters to recognize and bind multiple metabolites.

**Fig 5.**
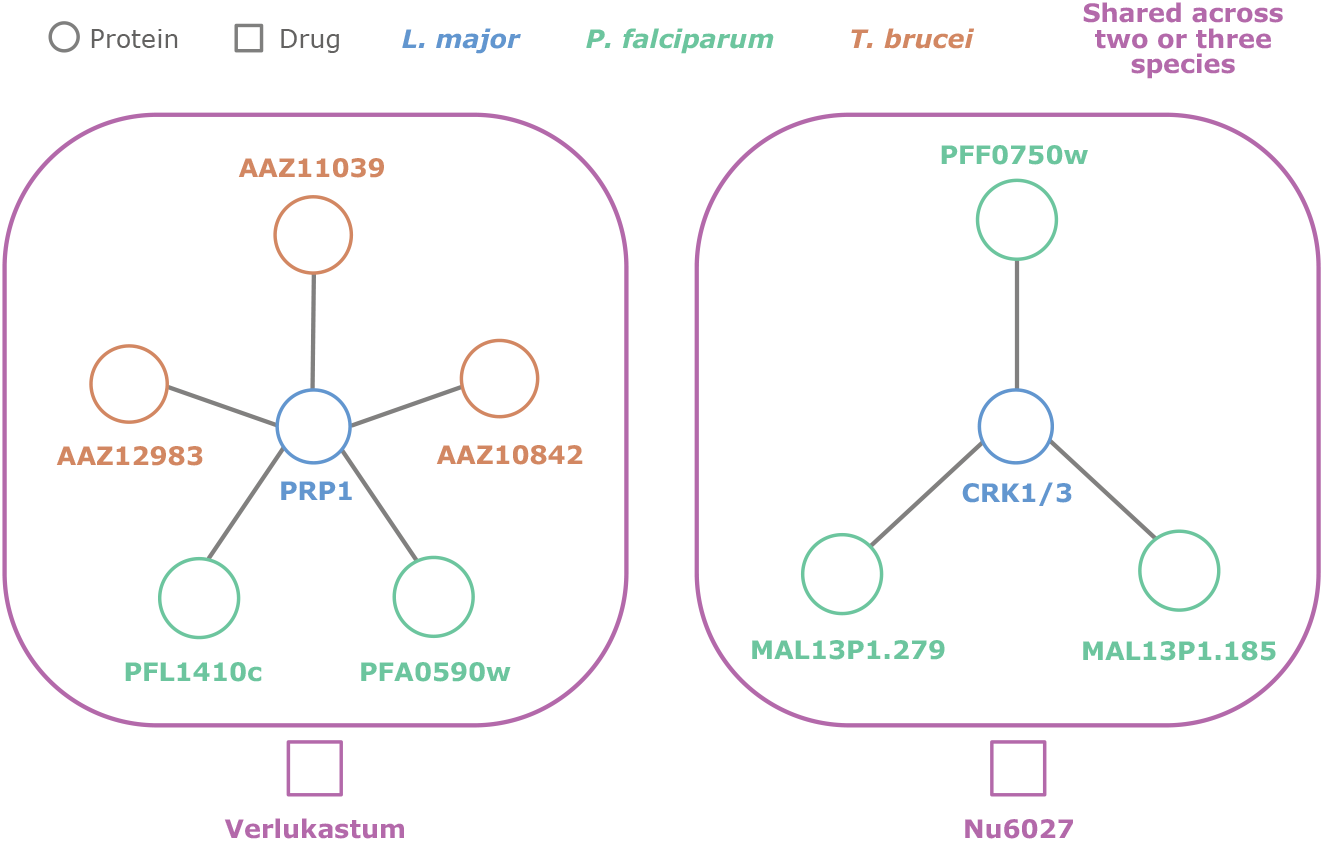
Drug-repurposing recommendations made by the final HGNN model. (Left) The drug verlukastum was predicted to interact with the pentamidine-resistance protein 1 (PRP1) in *L. major*. This protein is highly similar to five other proteins known to interact with verlukastum and that were seen during training. (Right) The experimental chemotherapeutic compound Nu6027 was predicted to interact with two *L. major* cdc2-related kinase proteins (CRK1 and CRK3). These two proteins are highly similar to three *P. falciparum* proteins seen during training and known to interact with Nu6027.

Together, these results point to novel approaches to expand the use of experimental drugs for treating leishmaniasis either as co-adjuvants (as in the case of verlukast) or as chemotherapy (as for Nu6027).

## Discussion and Conclusion

Despite the great advancements made in pharmaceutical chemistry and drug development, millions of patients still lack effective treatments against leishmaniasis and many other neglected tropical diseases. Without a strong economic incentive to justify large investments in research and development, it is unlikely that such treatments will ever get made under the traditional business model of the pharmaceutical industry. For that reason, it is critical to continue the development of drug repurposing techniques that may aid in the discovery of novel breakthrough therapies.

Here we have developed a drug repurposing model based on deep learning with hyperbolic graph neural networks. We leveraged drug-protein interaction data from taxonomically related organisms to train our model, and demonstrated its superior classification power over conventional machine learning and graph neural network models trained on the same data.

There are multiple examples in the literature with successful applications of graph theory and deep learning for drug discovery, repurposing, and predicting polypharmacy side effects [17–19, 39, 40]. However less studied is the use of hyperbolic graph neural networks in drug repurposing in general and in the context of leishmaniasis specifically. Since many of the previous studies in drug repurposing focus on human diseases and related pathways (e.g. cancer, type II diabetes) [3, 41, 42], we found it necessary to develop a multi-organism framework that leverages the knowledge from drug-protein interactions in other parasites. Therefore, this points to a potential future direction of our study in which we could consider the interaction of drugs with human proteins and their potential toxicity and off-target effects in the patient. For example, one could leverage predictions of drug side-effects and polypharmacy with models like Decagon [19], especially since our predictions around verlukast suggest a potential synergistic function with pentamidine.

A common criticism of computer-based drug design and discovery methods is that they often predict drugs that do not dock well to the target protein [43]. This is indeed a limitation of our study since we did not account for molecular docking in the drug-protein pairs predicted by our final model. This limitation stems from the lack of high-resolution structures of *L. major* proteins in the public domain and that were identified by our model in the inference set. However, a future direction of our study could address this limitation by making use of protein structures predicted by modern techniques such as AlphaFold2 [44] and RoseTTAFold [45]. These structures could then be used as input for predicting binding affinities of our predicted drugs with either traditional docking software [46] or more modern deep learning architectures such as MONN [47].

We also believe that the drug discovery can be aided by applying domain knowledge to identify specific protein interactions and pathways to improve modeling or evaluation [48]. A future direction can involve methods to assign higher scores to drugs that interact with subsets of important proteins rather than looking for high scores on individual or across all proteins in the testing set [49]. In conclusion, we have shown the potential of HGCNs for drug repurposing in the context of leishmaniasis. The code for this work is available here: https://github.com/layer6ai-labs/hgnn-dr. The framework developed in this work can be easily expanded to include other organisms and their drug-protein interactions, which we hope will contribute to the discovery of novel and safer treatments against leishmaniasis and other neglected diseases.

## Supporting Information

**Fig S1. Data splitting schematic** Diagram of the protein-drug interaction matrix illustrating the data splits into training, validation, test and inference sets. Here *P***_L_** and *D***_L_** refer to proteins and drugs with known interactions from the *L. major* organism network, and *P***_PT_** and *D***_PT_** refer to the same for *P. falciparum* and *T. brucei*.

**Table S1. HGNN and baseline model performance.** AUCPR, global top-*q*%, and local top-*k* metrics for model performance are reported for the test data split. The error reported is the standard deviation over ten random data split seeds.

## Appendix

The appendix contains additional details relevant for data splitting, feature extraction, model hyperparameters, and background on hyperbolic graph neural networks.

**Fig S1.**
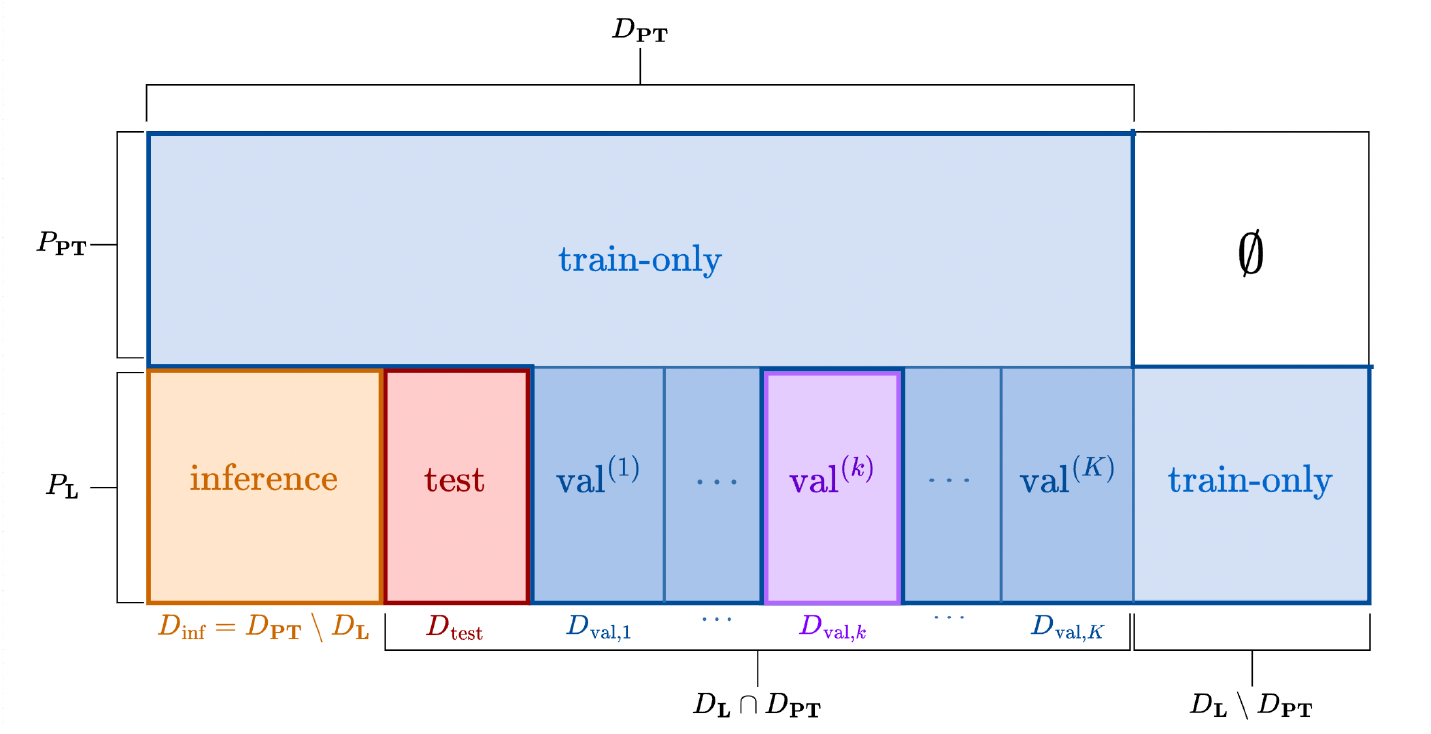
Detailed data splitting schematic. Diagram of the protein-drug interaction matrix illustrating the data splits into training, K-fold validation, test and inference sets. For a detailed description of the sets involved and the splitting procedure, see the Data Splitting section in the main text.

**Table S1.** HGNN-DR and baseline model performance. AUCPR, global top-q%, and local top-*k* metrics for model performance are reported for the test data split. The error reported is the standard deviation over ten random data split seeds.

| METRIC | LOGISTIC<br>REGRESSION | XGBOOST | LIGHTGBM | MLP | EGNN | HGNN-DR |
| --- | --- | --- | --- | --- | --- | --- |
| AUCPR | 0.037±0.017 | 0.434±0.056 | 0.314±0.051 | 0.214±0.071 | 0.544±0.026 | 0.659±0.029 |
| Top-0.5% Global Precision | 0.065±0.042 | 0.475±0.060 | 0.363±0.057 | 0.285±0.077 | 0.602±0.046 | 0.672±0.052 |
| Top-0.5% Global Recall | 0.058±0.031 | 0.433±0.071 | 0.331±0.058 | 0.259±0.048 | 0.547±0.017 | 0.612±0.036 |
| Top-1% Global Precision | 0.061±0.029 | 0.364±0.046 | 0.297±0.041 | 0.229±0.048 | 0.402±0.034 | 0.418±0.036 |
| Top-1% Global Recall | 0.110±0.041 | 0.662±0.077 | 0.541±0.080 | 0.416±0.052 | 0.730±0.033 | 0.759±0.037 |
| Top-3 Local Precision | 0.097±0.150 | 0.419±0.062 | 0.352±0.060 | 0.294±0.057 | 0.479±0.036 | 0.525±0.028 |
| Top-3 Local Recall | 0.133±0.210 | 0.482±0.040 | 0.412±0.064 | 0.330±0.073 | 0.482±0.048 | 0.555±0.038 |
| Top-5 Local Precision | 0.083±0.089 | 0.3559±0.058 | 0.305±0.056 | 0.252±0.056 | 0.402±0.032 | 0.437±0.032 |
| Top-5 Local Recall | 0.177±0.187 | 0.611±0.043 | 0.537±0.066 | 0.440±0.087 | 0.605±0.043 | 0.688±0.037 |

## Supplementary Text / Appendix

### Data splitting / K-Fold Validation

During training we randomly sample negative edges, in equal proportion to the number of positives. We use *K*-fold validation with *K* = 3 to perform model selection and hyper-parameter tuning. Once we determine the best model, we combine all of the validation data and train-only data to create a final model, and evaluate its performance on the test split. The protein-drug pairs in the inference set that are predicted to have a high likelihood to interact provide the final candidate drugs for targeting the *L. major* parasite.

### Protein and Drug Features

We use two chemoinformatics libraries, RDKit [26] and Mordred [25], for extracting molecular descriptors for each chemical in our network. Both of these packages compute numerical features of molecules based on their SMILES strings. Across all chemicals in the network, we remove descriptors that are either extremely sparse, highly skewed, or have constant values. The RDKit and Mordred libraries can compute up to 200 and 1,377 descriptors respectively. After removing features with a significant portion of zero or missing values, the number of descriptors is reduced to 140 and 1,155 respectively. We extract features for proteins using the UniRep model [27], which embeds amino acid sequences into 1,900-dimensional embedding vectors.

Features are not necessarily available for all proteins and drugs in our network. The FASTA sequence for a protein or SMILES string for a drug may not be available, and there are a small number of SMILES strings for which RDKit or Mordred descriptors can not be computed. However, the fraction of proteins and drugs for which features can be computed exceeds 99%. Missing features are imputed using their median values.

### Feature transformations

We use principal components analysis (PCA) with standard scaling to reduce the dimensionality of the Mordred and UniRep embedding vectors down to 100 each. PCA is applied to UniRep embeddings of all proteins across target and reference organisms, and similarly for Mordred descriptors across chemicals. Drugs or proteins with missing features are excluded from PCA.

We also apply the Yeo-Johnson power transform to the reduced features before passing them as input to the logistic regression, MLP, EGNN and HGNN-DR models. The performance of XGBoost and LightGBM does not improve by applying the Yeo-Johnson transform, so it is not applied for these models.

### Model Hyperparameters

#### Hyperparameter search

For feature-based models with early stopping, we set the maximum number of training iterations to 500, with 50 early stopping rounds. The iteration to use for the final models were determined by taking the maximum of the best iterations over 3-fold cross validation. For logistic regression, where this feature is not available, the maximum iterations is tuned as a hyperparameter for grid search.

For each feature-based model, we obtain final hyperparameters by performing randomized grid search using 3-fold cross validation: For logistic regression, we search over `max_iter` ∈ {200, 300, 400}, as well as the L2 regularization strength C ∈ {0.01, 0.1, 1.0, 10, 100}. For XGBoost, we search over `max_depth` ∈ {6, 10, 14}, learning rate `eta` ∈ {0.01, 0.05, 0.1, 0.5, 1}, as well as L1 / L2 regularization strength ∈ { } ∈ { } − ∈ { } ∈ { } ∈ { } `alpha, lambda` 0, 0.01, 0.1, 1, 10. For LightGBM, we search over the same grid, with the inclusion of `max_depth` = 1 (unlimited). Finally, we search over MLP architectures with `hidden_layers` 1, …, 5, each having `hidden_dim` 20, 40, 60, and each hidden layer is followed by a ReLU nonlinearity. We use the Adam optimizer with no batching, learning rate lr 0.001, 0.01, 0.1, 1, and `weight_decay` 0, 0.01, 0.1, 1, 10. Hyperparameters for GNN models were manually tuned.

#### Final hyperparameters

For logistic regression, we use: `max_iter` = 200 and C = 100. For XGBoost: `num_boost_round` = 433, `lambda` = 10, `alpha` = 0, `eta` = 0.1, and `max_depth` = 10. For LightGBM: `num_boost_round` = 392, `lambda_l1` = 0.1, `lambda_l2` = 10, `learning_rate` = 0.5, and `max_depth` = 10. Finally, for MLP we use a hidden dimension of 40, with 3 layers and weight decay of 0.01 and a learning rate of 10*^−3^*. For the Euclidean GNN we use a 2 layer encoder, and HGNN-DR is trained with 2 layers of tangent aggregation. We use tanh activation and an embedding dimension of 25 with a learning rate of 10*^−^*2 and Adam optimizer.

### Hyperbolic Graph Neural Networks Background

In this section we provide background on hyperbolic space relevant to the formulation of HGNNs. Hyperbolic space is a Riemannian manifold with constant negative curvature. There are many equivalent formulations of hyperbolic space. In this work we use the Poincaré Ball model with curvature *c* = *−*1. It is defined by the open sphere, *ℬ^n^*, with the metric tensor, *g_B_*, as follows: *ℬ^n^* = {***x*** ∈ *ℝ^n^* : ||**x**|| *<* 1} and *g_B_* = (2*/*(1 *−*||**x**||^2^))^2^𝕀*_n_*, where ||. denotes the Euclidean norm.

The tangent space to a point **x** *∈ ℬ^n^*, 𝒯**_x_***ℬ^n^*, is a Euclidean space that approximates *ℬ^n^* at **x**. The exponential map exp**_x_** : *𝒯***_x_***ℬ^n^* → *ℬ^n^* and logarithmic map log**_x_** : ℬ*^n^ → 𝒯***_x_***ℬ^n^* allow points in the tangent space to be mapped to hyperbolic space and visa-versa. For points **x**, **y** *∈ ℬ^n^* and **v** *∈ 𝒯***_x_***ℬ^n^*, they are defined as [34],

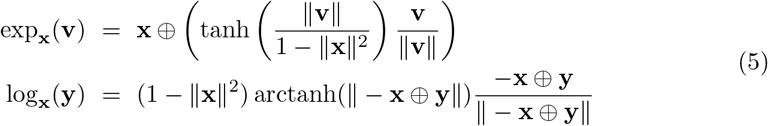

where **y** ≠ **x**,**v** ≠ 0 and the Möbius addition *?* is defined as

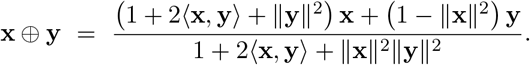

Hyperbolic feature transformation utilizes the use Möbius matrix multiplication operator, ⊗, defined as [34]

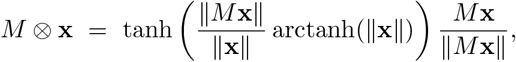

if *M* **x** *≠* 0 and *M ⊗* **x** = 0 otherwise.

In the Poincaré Ball model the distance between two points **x**, **y** *∈ ℬ^d^* is given by

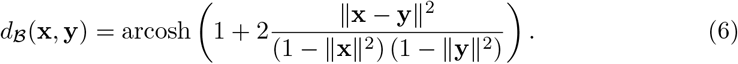

For a fixed point **x**, the distance to **y** grows exponentially as **y** approaches the boundary.

